# Enabling the execution of large scale workflows for molecular dynamics simulations

**DOI:** 10.1101/2021.04.14.439795

**Authors:** Pau Andrio, Adam Hospital, Cristian Ramon-Cortes, Javier Conejero, Daniele Lezzi, Jorge Ejarque, Josep LL. Gelpi, Rosa M. Badia

## Abstract

The usage of workflows has led to progress in many fields of science, where the need to process large amounts of data is coupled with difficulty in accessing and efficiently using High Performance Computing platforms. On the one hand, scientists are focused on their problem and concerned with how to process their data. On top of that, the applications typically have different parts and use different tools for each part, thus complicating the distribution and the reproducibility of the simulations. On the other hand, computer scientists concentrate on how to develop frameworks for the deployment of workflows on HPC or HTC resources; often providing separate solutions for the computational aspects and the data analytic ones.

In this paper we present an approach to support biomolecular researchers in the development of complex workflows that i) allow them to compose pipelines of individual simulations built from different tools and interconnected by data dependencies, ii) run them seamlessly on different computational platforms, and iii) scale them up to the large number of cores provided by modern supercomputing infrastructures. Our approach is based on the orchestration of computational building blocks for Molecular Dynamics simulations through an efficient workflow management system that has already been adopted in many scientific fields to run applications on multitudes of computing backends.

Results demonstrate the validity of the proposed solution through the execution of massively parallel runs in a supercomputer facility.

## I. Introduction

Computational workflows are one of the most used tools to assemble and run simulations of different scientific fields as climate predictions, bioinformatics, engineering, etc. Researchers can compose their applications, usually made of pieces of code available in libraries and binaries, using a textual or graphical representation of the dependencies between those parts, and let the runtime of the workflow management system to orchestrate the execution on a given computational platform. In particular, HPC systems are getting more and more attractive for the run of workflows that have been traditionally executed on distributed systems as grids or clouds, and that can have tasks that require a certain degree of parallelism (i.e., MPI tasks). In general, the trend is to have complex HPC systems built on hybrid architectures that include traditional processors with accelerators, and that can be used together with cloud infrastructures. On top of the computing complexity, the packaging of workflows is an additional issue, with containers becoming a popular way to distribute and deploy applications.

To address the issues above, it is a must to have a workflow management system that can efficiently orchestrate the applications on modern computing infrastructures, that can offer a simple interface for the composition of the tasks, and that provide advanced capabilities such as task level fault-tolerance and automatic resource reservation for different kind of loads including MPI. This paper presents the proposal of a framework for the definition and orchestration of biomolecular simulations on HPC infrastructures, that satisfies the above mentioned requirements. The building blocks software library developed by the BioExcel Centre of Excellence has been used to implement pipelines that automatize the various steps of Molecular Dynamics simulations that are in many cases performed manually by the users. The PyCOMPSs programming framework has been adopted to define the tasks of the workflows and to parallelize their execution. The definition of the tasks allows to automatically reserve the proper number of computing nodes and cores taking into account specific requirements and constraints of each task, and to execute the internal tools (i.e., MPI) transparently to the user. Furthermore, the developed building blocks are proposed on multiple alternative choices for deployment including HPC, Clouds, containerized platforms or environment management systems as Conda. The workflow has been run in the Marenostrum IV supercomputer using 38,400 cores in parallel.

The paper is structured as follows: Section II describes the building blocks software library considered as a target use case for the large scale execution of biomolecular workflows, and Section III details the design and development of the workflows used to validate the proposal using PyCOMPSs. Next, Section IV reports the results of the execution of the developed workflow on an HPC infrastructure. Section V presents the state of the art and related work on topics involved in the proposed research. Finally, Section VI wraps up the paper.

## II. Application building blocks for computational biomolecular simulations

BioExcel Centre of Excellence for computational biomolecular research (BioExcel CoE) is a European initiative whose primary purpose is to give support to academic and industrial researchers in the use of high-performance and high-throughput computing (HPC & HTC). BioExcel goals include i) to optimize and increase the scalability of biomolecular simulations using the principal codes involved in the project: GROMACS [1] (MD), HADDOCK [2] (Docking), and CP2K [3] (QM/MM); and ii) ease the usability of biomolecular computational tools and biological databases through a range of scientific workflows and associated deployment environments. Linking both goals together is of great interest for the field since it gives the opportunity to easily build complex scientific workflow systems that benefit from the inherent scalability of the biomolecular simulation tools.

However, joining different biomolecular tools in a complex pipeline is not always straightforward. Interoperability between tools is a complex issue in biomolecular simulations, mainly due to the lack of a set of standards. BioExcel building blocks (biobbs) [4] is a software architecture designed to tackle the interoperability problem, thanks to a simple wrapping approximation. Biobbs are a collection of small wrappers written in Python and organized in layers. Each building block encapsulates software components (1st layer) and provides a well-defined interface for input, output, configuration, and provenance (2nd layer). A standardized syntax is used in all of the building blocks, with each of the wrappers internally performing the necessary format conversions for input and output, and launching the tool inside, which runs unaltered. With this design, a large set of biomolecular tools can be launched using a homogeneous syntax, providing as well a uniform and stable interface with enough information to plug the components into interoperable workflows. The optional 3rd layer is used to adapt the building blocks to different workflow managers, from Graphical User Interfaces such as Galaxy or KNIME, to HPC-designed tools such as Toil or PyCOMPSs. In Section III-C we explain how we have implemented PyMDSetup with PyCOMPSs to parallelize the execution of the building blocks.

## III. Parallelization design

Before going into the parallelization design, we introduce PyMDSetup and PyCOMPSs; respectively, the target application and the framework used to parallelize it. Next, we focus on the mechanisms that PyCOMPSs provides to exploit the maximum performance considering its requirements and how they have been used.

### A. PyMDSetup and biobb_wf_mutations

PyMDSetup is an automated protocol to model residue mutations in 3D protein structures detected from genomics data, and prepare and run Molecular Dynamics simulations for all the generated structures. The pipeline receives a PDB file (wild type protein 3D structure) and a set of mutations as input. Next, it prepares and runs MD simulations for each of the systems, thus obtaining static information (an ensemble of modelled structures for each of the protein variants), and dynamic data (trajectories for each of the protein variants). Both types of information can later be used in a comparative study.

The complete PyMDSetup workflow is represented in Figure 1. It starts with a simple loop around the input collection of protein variants, with each loop modelling the corresponding variant, preparing the structure for a MD simulation (MD setup), and finally running the simulation. All the MD steps are run using the GROMACS MD package. The first step is modelling the mutated variant of the input protein, using the SCRWL4 program [5]. In the next step, a GROMACS topology is created, defining the modeled structure in terms of geometrical properties, spatial relations and force field parameters. The following two steps build a system box surrounding the protein and fill it with water molecules, mimicking the hydrated environment where proteins perform its function in the living cells. Some of the water molecules added in the system box are replaced by monoatomic ions in the next steps, to neutralize the energy of the system, which is needed when working with periodic boundary conditions (PBC) for an accurate and efficient treatment of electrostatic interactions. After that, a determined ionic concentration of 0.05M is added to mimic the physiological ionic strength. Next steps of the pipeline correspond to an extended MD setup pipeline needed in complex or big systems, composed of two energetic minimization steps and eight system equilibration steps. Details on the simulation configuration (force field, ensemble, setup parameters, minimization and equilibration restraints and time) are described elsewhere [4]. Finally, with the system completely equilibrated, a 5ns-length free (unbiased) MD simulation is launched as a final step in the pipeline.

**Fig. 1.**
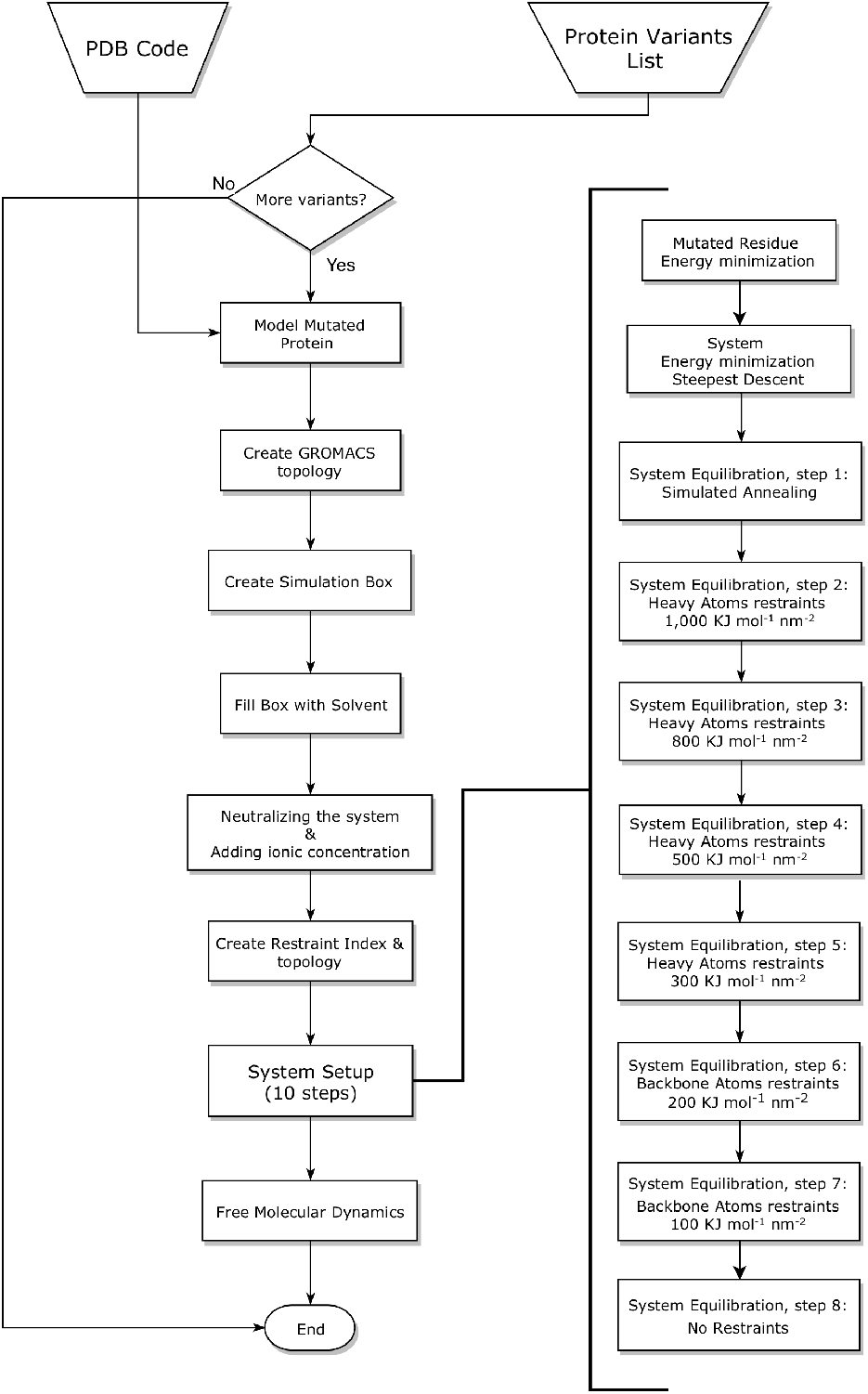
PyMDSetup flowchart

The test case chosen to run a massively parallel execution using the described workflow is the Pyruvate Kinase protein (PDB code 2VGB [6]). Pyruvate Kinase is a widely studied enzyme in biochemistry, due to its major role in the regulation of glycolysis. It catalyzes the irreversible conversion of phosphoenolpyruvate (PEP) to pyruvate, generating an ATP molecule in the process. More specifically, the structure used is a representative of the human erythrocyte pyruvate kinase (R-PYK), one of the four known human pyruvate kinase isoforms, encoded by gene PKLR and expressed in erythrocytes. It has been selected because of its large number of annotated missense variants that have been associated to a disease called non-spherocytic hemolytic anemia, a rare, autosomal recessive disease that causes blood disorders characterized by the premature destruction of red blood cells (erythrocytes), resulting in anemia. To be able to get the most out of the combination of PyCOMPSs and the PyMDSetup workflow, a set of 200 mutations consisting on reported pathogenic variants were manually selected from the whole set of variants available at the UniprotKB database [7], and used as input. With these inputs, the generated workflow can sequentially run 200 totally independent MD simulations, which are then parallelized by PyCOMPSs.

A shortened, simplified version of the PyMDSetup workflow called biobb_wf_mutations was used to run the validation tests (see Section IV). This workflow, generated initially to present the BioExcel building blocks to the computational biomolecular community, is basically performing the same work than the original one: looping around a collection of protein variants, modelling the mutation and computing an MD simulation from each of them. The difference relies on the MD setup process (Figure 1), which in this case is reduced from 10 steps to just three steps. These three steps consist of an energy minimization followed by two steps of equilibration: the first one with constant volume (NVT ensemble), and the second one with constant pressure (NPT ensemble). This approach is based on the well-known GROMACS MD setup tutorial [8], which was built with teaching purposes and is only valid for molecules with small complexity. When working with complex systems, such as the one recently introduced, more elaborated approximations such as the PyMDSetup pipeline are recommended.

### B. PyCOMPSs

PyCOMPSs is a task-based programming model that enables the parallel execution of sequential Python code with minimal effort. To this end, it provides a set of Python decorators that allow the user to identify the function/methods to be considered as tasks and a small API for synchronization. PyCOMPSs also features a runtime that is able to identify the data dependencies that exist among tasks and to extract the parallelism between them building a data dependency graph of tasks. The runtime is also responsible of managing their execution across distributed infrastructures (e.g., Grids, Clusters, Clouds, and Container manager clusters) – scheduling them and performing the necessary data transfers when needed – guaranteeing that the result is the same as if the application was executed sequentially.

The main decorator that PyCOMPSs provides to identify that a function/method has to be considered as a task is the *@task* decorator. This decorator can be placed on top of any function, instance method or class method and it is used to identify the function’s input/output parameters and return peculiarities.

Moreover, PyCOMPSs provides a set of decorators to identify that the execution of a binary is considered as a task. In particular, PyCOMPSs support the invocation of three types of binaries: simple binaries, MPI binaries, and OMPSs. To this end, the *@binary, @mpi*, and *@ompss* decorators respectively have to be placed on top of the *@task* decorator. The users must also specify the binary to execute, as well as the specific parameters for MPI and OmpSs invocation (i.e., the number of nodes to use). The *@task* and *@mpi* have been used to parallelize the PyMDSetup application. For this reason, they will be described with further detail in the next subsection.

Besides, PyCOMPSs also supports the tasks constraint definition. To this end, it provides the *@constraint* decorator, which also needs to be placed on top of the stack of decorators. Constraints are also used to let the developers provide hints on the fault tolerance at task level thus allowing to discard parts of a workflow that don’t lead to relevant results or that fail for some reason, without affecting the main application. In the related work we provide more details on the reasons why PyCOMPSs is a better solution for this kind of applications.

### C. Applying PyCOMPSs to PyMDSetup

The first step was to identify in the PyMDSetup code the potential functions to be considered as tasks, as well as the ones that could benefit from a particular PyCOMPSs specific decorator.

Since the PyMDSetup was sorted out into functions per step, we considered each step as a task. The two main reasons for this decision were: 1) to allow to modify the mutation analysis, including new steps, doing other or repeating already existing steps, and sort them; and 2) since PyMDSetup worked sequentially and each step included the necessary invocations to binaries with parameter parsing and pre/post processing, they represented perfectly the unit of work for a task. We considered the idea of using the *@binary* decorator, but it was discarded since each step does specific parameter parsing and includes specific processing using Python.

For example, the *scwrl* function (see Listing 1) generates an `scwrl.Scwrl` object, and calls its `launch` method, which contains the code to perform the *Scwrl* step. This function consumes a file (provided through the `input_pdb_path`) and produces another as a result (stored in the path indicated by the `output_pdb_path` parameter). This function has been decorated with the *@task* decorator, and defined that the parameter `input_pdb_path` is a FILE with IN direction, while `output_pdb_path` is also a FILE, but with OUT direction.

**Listing 1:**
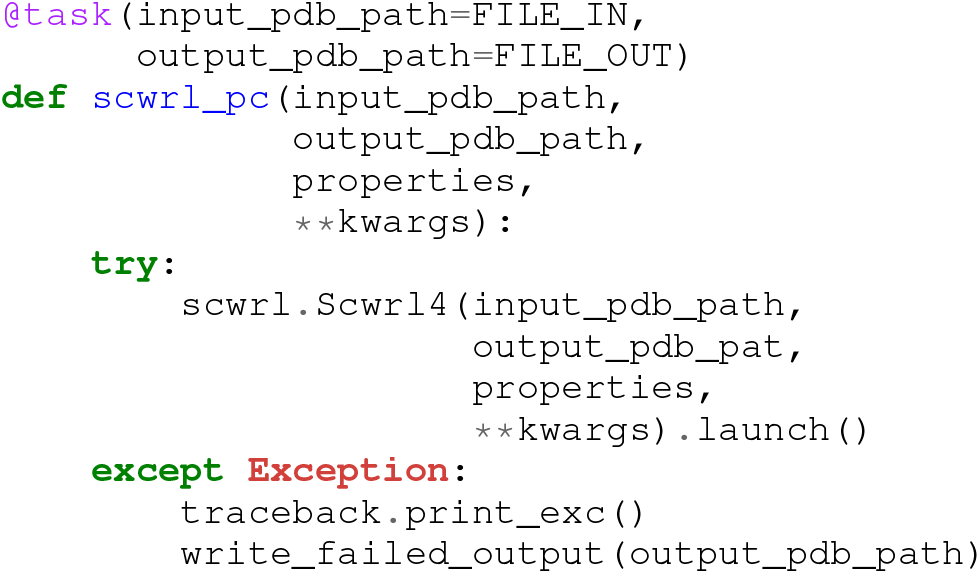
@task decorator example.

However, there where three steps (*mdrun_pc, mdrun_pc_cpt* and *mdrun_pc_all*) that internally execute Gromacs with different parameters. Since they did not require parameter parsing nor specific Python code, and Gromacs can benefit from running internally in parallel with MPI, we decided to use the *@mpi* decorator in conjunction with the *@constraint* decorator.

An example of this usage is shown in Listing 2. It depicts the *mdrun_pc* function, which invokes gromacs with the `mdrun` parameter, and then, consumes an input file and produces a set of output files (each one preceded by its required flag). It has been decorated with *@mpi*, specified the MPI runner and the binary to invoke (`gmx_mpi`), and the number of nodes that will be assigned to its execution (`computing_nodes`). Thus, it has been decorated with *@constraint* to define the number of processes that will be spawned per computing node (`computing_units`).

**Listing 2:**
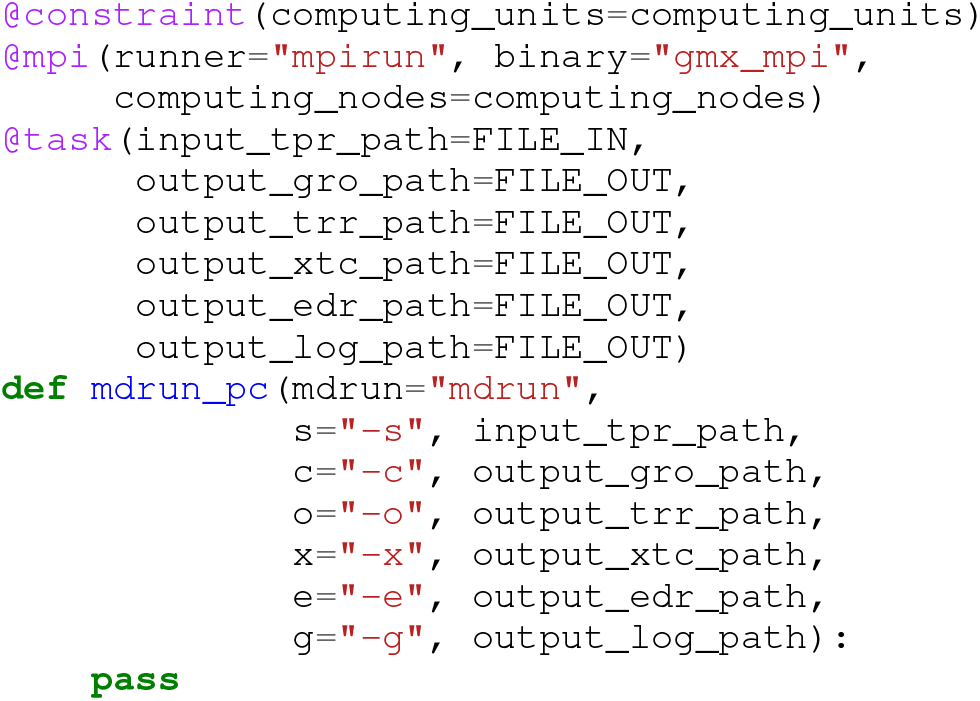
@mpi decorator example.

When the *mdrun_pc* function is invoked, the runtime runs `gmx_mpi` in a worker node considering all defined task parameters, as shown in Listing 3. The call provides to the *mpirun* the number of nodes and computing units, as well as the file with the hostnames where to run the MPI binary, and all binary arguments in the same order as defined in the task function.

**Listing 3:**
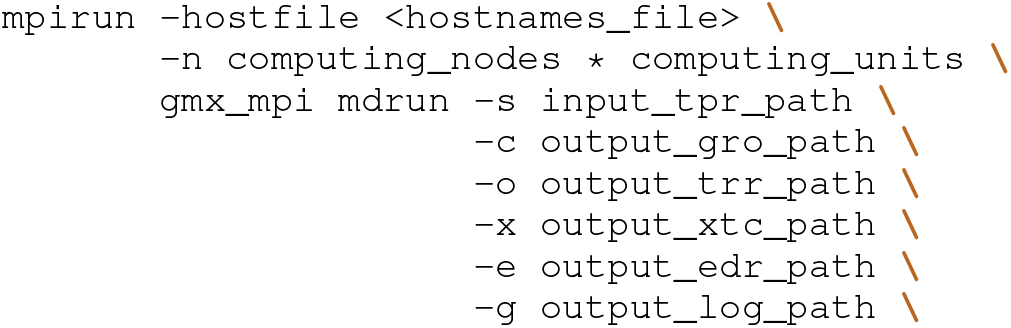
mdrun_pc invokation equivalence.

Finally, we included a *compss_barrier()* after the main mutations loop, so that the execution does not continue until all mutation analysis finish.

### D. Parallelization analysis

The main benefit observed from the application of the parallelization described in the previous section is that the process of analysing a mutation can be performed in parallel with any other mutation, enabling to examine multiple mutations at the same time (if enough resources are available). Besides, the COMPSs runtime can detect some parallelism within each mutation. This can be easily observed in the task dependency graph that the COMPSs runtime builds (Figure 2), which shows a two mutation analysis.

**Fig. 2.**
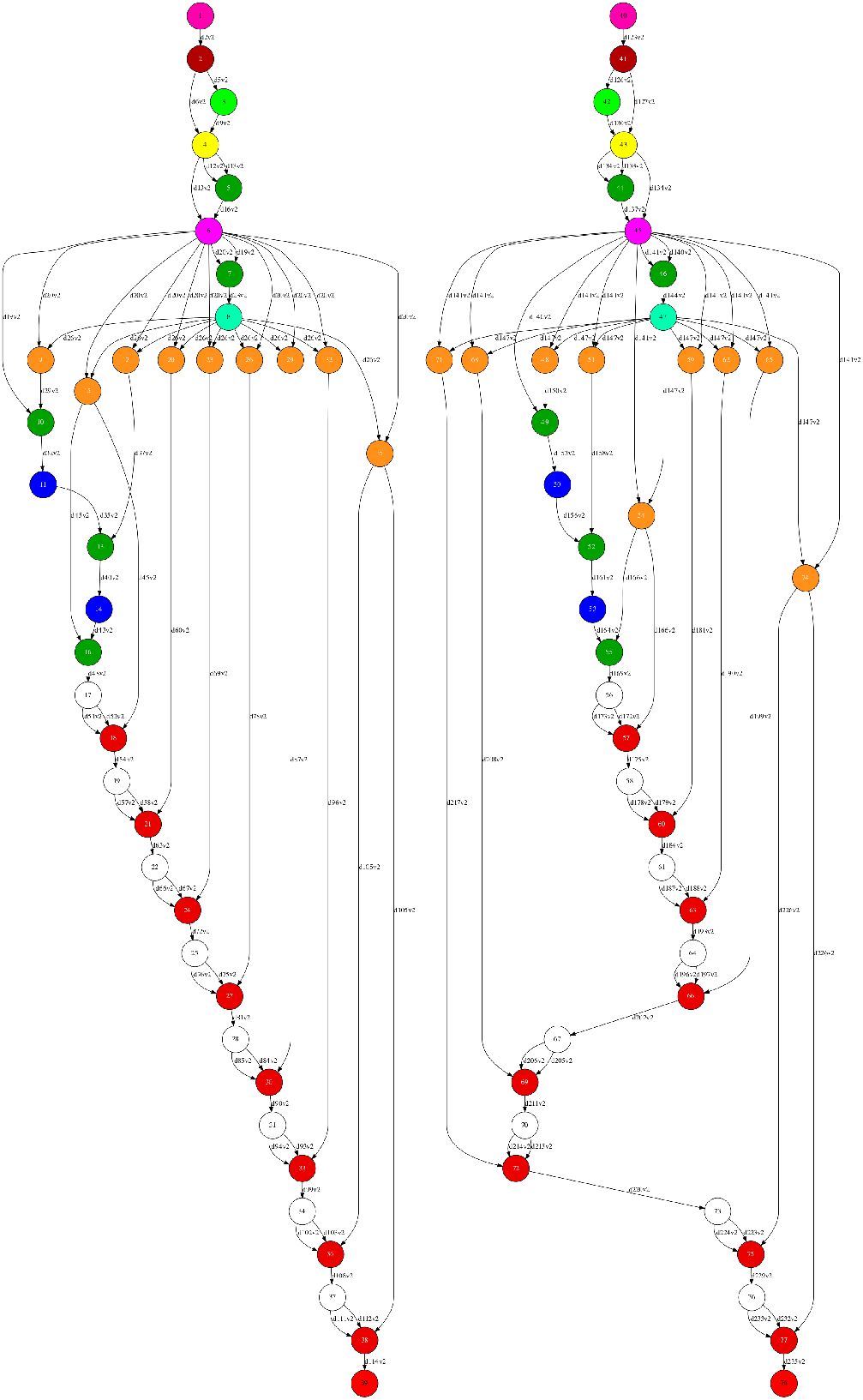
Task dependency graph (2 mutations)

In particular, there are 12 defined tasks in PyMDSetup, and each mutation requires 39 invocations (39 tasks for mutation). Moreover, the PyMDSetup also includes two extra tasks to perform a reduction of the results and generate a plot. The plot can be deactivated since it becomes useless when performing the analysis of lots of mutations (e.g., more than 16). In this case, a post-mortem analysis of the files produced by the execution becomes necessary.

Concerning the code, Table I shows the PyMDSetup lines of code, remarking the PyCOMPSs lines which enable this parallelization. Notice that with only 31 lines of Python code, the parallelism of the whole application can be achieved without dealing with any of the complexities of parallelization. The PyCOMPSs impact on the code represents less than 1.5%, just composed by 20 decorators (14 *@task*, 3 *@mpi* and *3@constraint*), 6 imports, and 1 *compss_barrier*. Finally, the lines required in shell scripts (Bash) are related to the run/launch commands provided by PyCOMPSs.

**TABLE I.** Lines of code

| Language | Files | Blank | Comment | Code | PyCOMPSs lines |
| --- | --- | --- | --- | --- | --- |
| Python | 22 | 314 | 333 | 1839 | 27 |
| YAML | 2 | 94 | 112 | 936 | 0 |
| Markdown | 1 | 27 | 0 | 95 | 0 |
| Bash | 9 | 14 | 60 | 87 | 4 |
| Total | 34 | 449 | 505 | 2967 | 31 |

In conclusion, we have been able to parallelize the PyMDSetup application with PyCOMPSs with a minimal impact on the code, achieving relevant parallelism, and hiding the parallelization complexities to the developer and user.

## IV. Validation

We have performed a set of experiments to validate the efficiency of PyCOMPSs parallelizing the PyMDSetup workflow to scale to large supercomputing infrastructures. To perform these experiments, we have set up two MD analysis with the parallelised version of PyMDSetup:

1. Reduced PyMDSetup (biobb_wf_mutations): A 16-step MD analysis to evaluate how the different application parameters and resource configurations affect the execution scalability, and how the PyCOMPSs parallelization can be tuned to get the maximum execution performance. The code of this experiment is available at [9].
2. Complete PyMDSetup (PyMDSetup): The study of the Pyruvate Kinase protein where we evaluate the massive parallelization of the PyMDSetup tool when we study a large number of mutations using thousands of CPUs in parallel. The code of this experiment is available at [10]

### A. Infrastructure

The results presented in this section have been obtained using the MareNostrum IV Supercomputer [11] located at the Barcelona Supercomputing Center (BSC). Its current peak performance is 11.15 Petaflops, ten times more than its previous version, MareNostrum III. The supercomputer is composed by 3,456 nodes, each of them with two Intel^®^Xeon Platinum 8160 (24 cores at 2,1 GHz each). It has 384.75 TB of main memory, 100Gb Intel^®^Omni-Path Full-Fat Tree Interconnection, and 14 PB of shared disk storage managed by the Global Parallel File System.

### B. Results

#### 1) Reduced PyMDSetup (biobb_wf_mutations)

This section presents the results of the experiments performed with the reduced workflow with the parallelized version of PyMDSetup. In this section, we have conducted several runs with different application parameters and several resource configurations to know how the application performs depending on these parameters. The first experiment aims at evaluating the performance of a single mutation analysis depending on the analysis problem size and on the number of computing nodes assigned to each part of the simulation. The duration of this 16-step MD analysis is dominated by the *mdrun* tasks which run a Molecular Dynamics simulation using the GROMACS software. The duration of this simulation grows with the number of simulation steps and can be parallelized with MPI.

Figure 3 shows the scalability of a single mutation analysis for different simulation steps when increasing the number of nodes per *mdrun* task from one to eight. In each node, we spawn 48 MPI processes. Hence, the simulation uses from 48 to 384 MPI processes. This processes-to-task mapping has been easily configured modifying the `computing_nodes` and `computing_units` properties of the PyCOMPSs decorators. The figure shows that for higher (1M5Steps) simulation steps the scalability is better and, up to 4 nodes, it is close to ideal. However, with more than 4 nodes, the scalability starts to worsen.

**Fig. 3.**
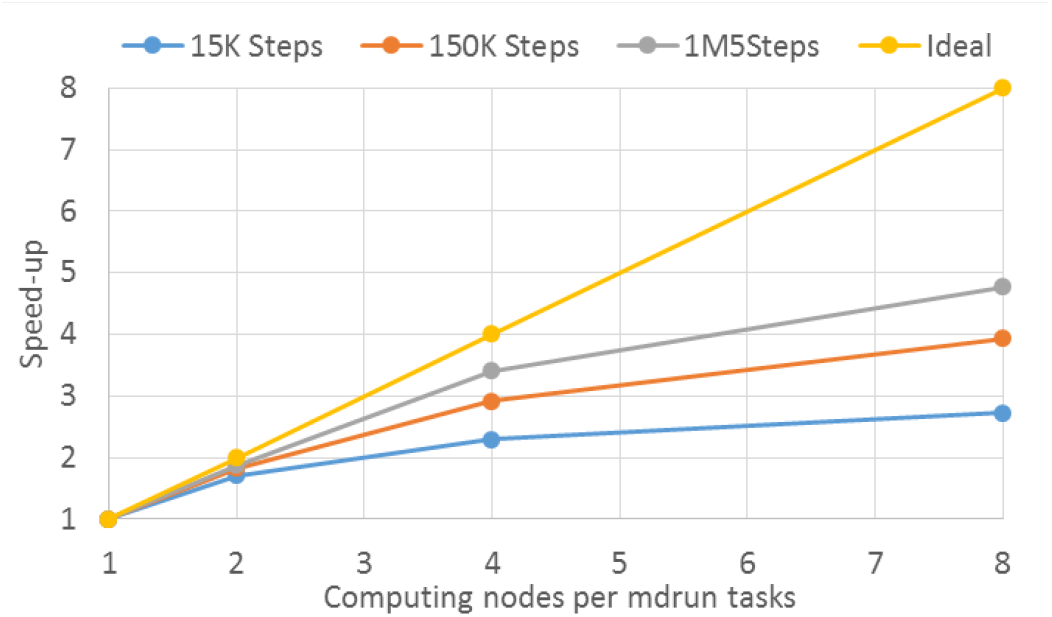
Scalability of one mutation analysis using different computing nodes and simulation steps for the *mdrun* task executions

The second experiment focuses on evaluating the scalability of the PyCOMPSs parallelization by performing a strong and weak scaling analysis. Figure 4 shows the results of the strong scaling analysis. In this case, we have evaluated 48 mutations, 15K simulation steps per evaluation and mapping one node (48 MPI processes) per *mdrun* task. The figure shows that scalability is growing close to the ideal. The results of the weak-scaling analysis are shown in Figure 5, where we have increased the number of computing nodes with the number of mutations. In this case, we have seen that the time is quite similar in all the runs, and the efficiency is higher than 90% in all cases.

**Fig. 4.**
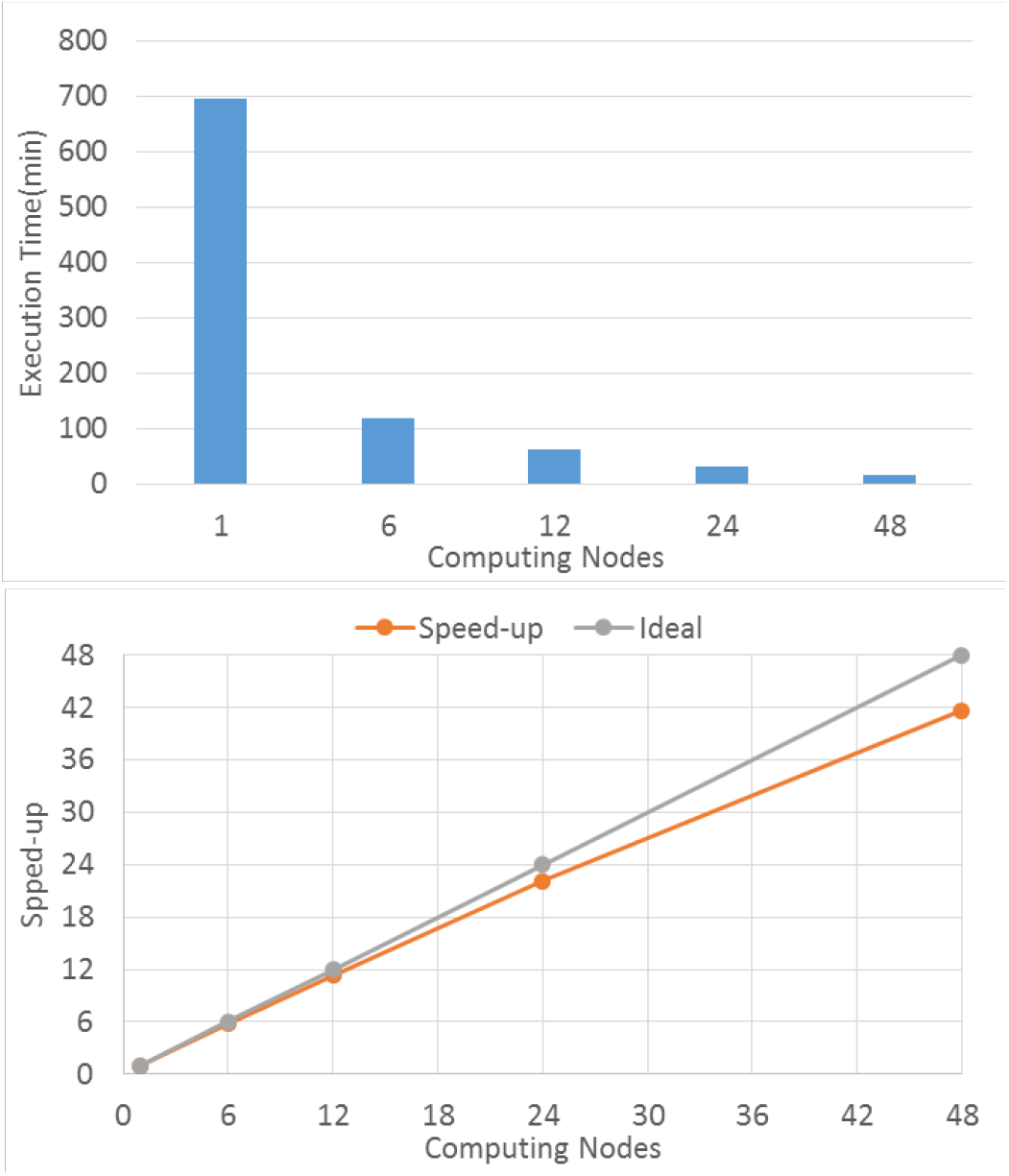
Strong scaling results for 48 mutations analysis (one node per *mdrun* and 15K steps analysis using different computing nodes for the *mdrun* task executions

**Fig. 5.**
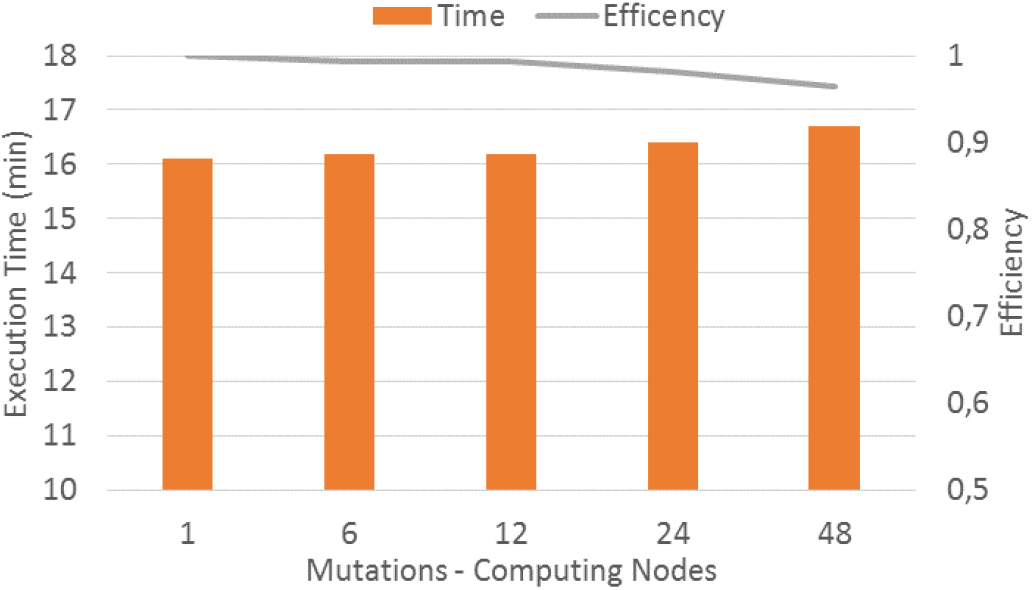
Weak scaling results with a load of one mutation analysis of 15K steps

Analysing the previous results (mdrun task parallelism and the whole workflow parallelism), we can observe that there is a trade-off between the number of parallel tasks and the grade of parallelism inside the *mdrun* tasks. For a fixed number of resources per application, the more resources are assigned to tasks the less parallel tasks are run. However, in this case, having more tasks in parallel leads to a better scalability than using more resources per task. Therefore, we can conclude that the optimal setup is assigning the resources per task to achieve the maximum number of tasks in parallel. Figure 6 depicts the mentioned trade-off. It shows the execution time of a different number of mutations and the *mdrun* task configuration, using a total of 24 computing nodes. The results confirm our conclusion: with 24 mutations, we achieve the best time using one node per task; with 12 mutations, we achieve the best time using two nodes per tasks, and so on.

**Fig. 6.**
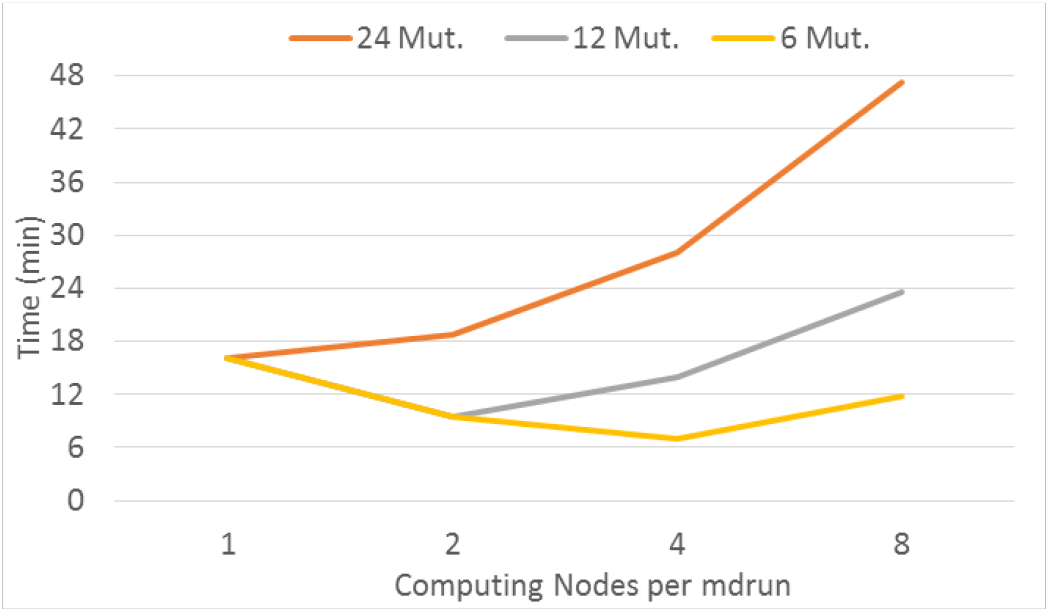
Mutations/nodes-per-mdrun trade-off (24 computing nodes)

Notice that setting this optimal configuration with Py-COMPSs is very easy since the users can pre-calculate the number of nodes per task by comparing the number of mutations and nodes, and assign it to the `computing_nodes` property of the PyCOMPSs decorator.

#### 2) Complete PyMDSetup

This section focuses on the analysis, using the Paraver [12] tool, of massively parallel execution of the workflow on 800 nodes of MareNostrum IV (which involves 38,400 cores) with the Pyruvate Kinase protein. Paraver is a powerful performance visualisation and analysis tool based on traces developed at BSC that can be used to analyse any information that is expressed on its input trace format. In the case of PyCOMPSs applications, its runtime is instrumented with Extrae [13], a BSC instrumentation package that can generate trace files for Paraver. Producing trace files for PyCOMPSs users is enabled by setting an execution flag.

The total number of tasks executed in this workflow instance is 7,800. Three main tasks for each mutation dominate the workflow execution: pyruvateKinase_MN.mdrun_pc (performs a minimization process), pyruvateKinase_MN.mdrun_pc_cpt (implements the equilibration of the data) and pyruvateKinase_MN.mdrun_pc_all (executes the final simulation). These three tasks are all parallel invocations (with MPI) of GROMACS executed in 4 nodes. Figure 7 shows a timeline with the task view of this execution, where each colour represents a task type.

**Fig. 7.**
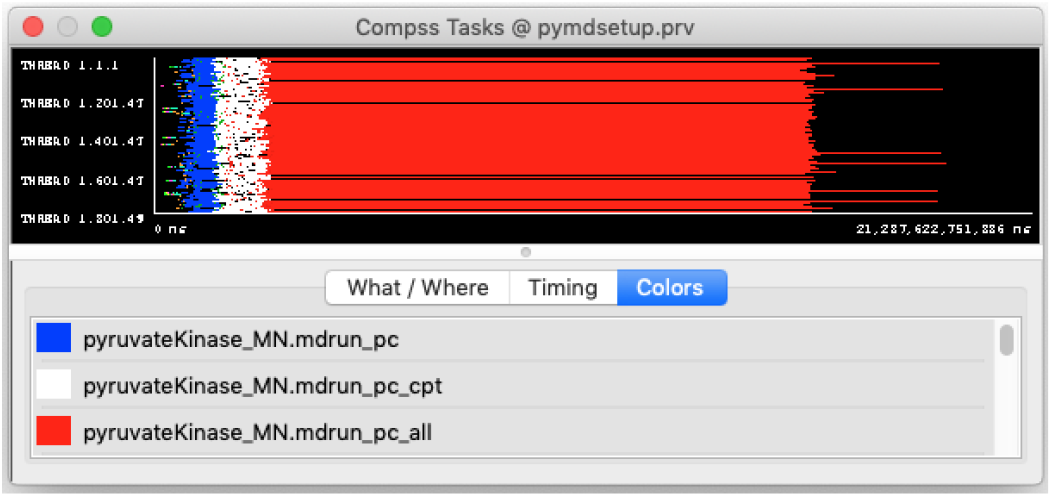
Paraver timeline view of the tasks of the Pyruvate Kinase workflow

The 7,800 tasks are generated at the beginning, and their dependencies are analysed. For each of the tasks, the runtime takes an average of 709.75*μs* per task to register it, an average of 1.80*ms* per task to analyse its dependencies with other tasks, and an average of 4.80*ms* to schedule the task if it is ready for execution (the task does not have dependencies with previous tasks). Every time a task finishes, the runtime updates the task graph (average of 573.30*μs*), and if new tasks are now ready for execution, it tries to schedule them to available resources (average of 120.39*ms*).

These processes are performed by three different threads in the runtime and partially overlapped between them in such a way that these overheads do not directly impact the tasks’ spawning time. Figure 8 shows a Paraver capture of the beginning of the application execution highlighting the behaviour of the three runtime threads. The different colours represent the different actions performed by the runtime.

**Fig. 8.**
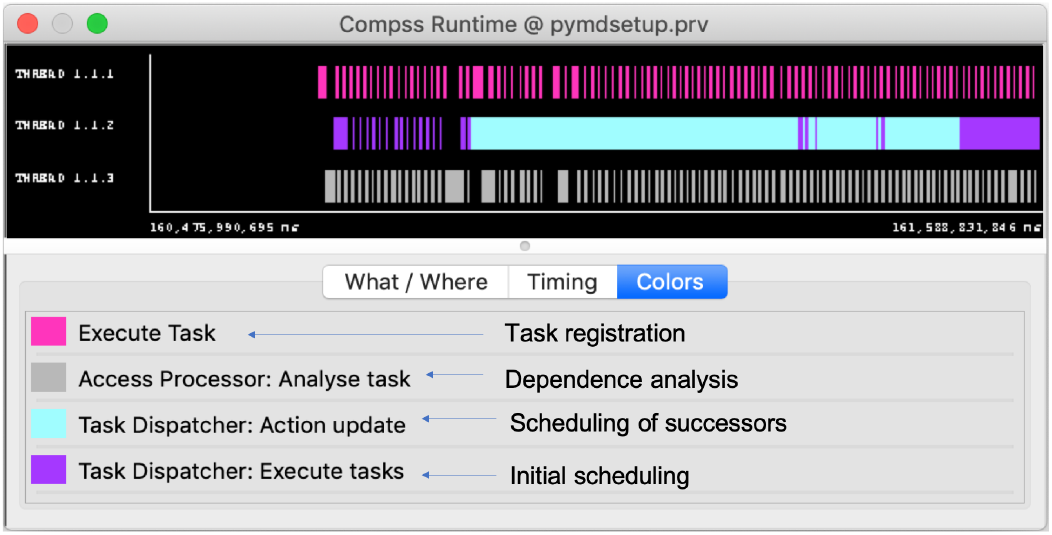
Paraver timeline view of the runtime behaviour

The interval between the registration of the first task and its execution is of 520.71*ms*. From there, the interval to schedule a new sequential task takes between 375 – 429*ms*. However, when the task to be scheduled is a multi-node one (GROMACS MPI simulation), this time grows up to 1, 750*ms*. In any case, since the MPI simulations dominate the execution, once the tasks have been scheduled, the computation resources are all used. Figure 9 shows the profile of the number of tasks on execution at each moment of the run, being the top value the 800 × 48 cores of the allocation. This figure shows that for most of the execution time, the allocation is 100% used.

**Fig. 9.**
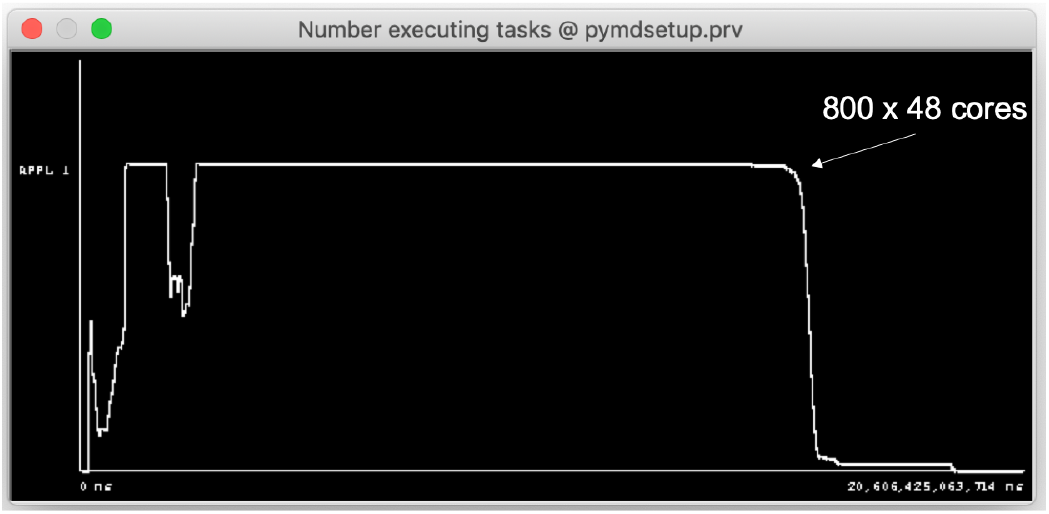
Paraver view showing the number of tasks on execution at each moment

We have also analysed the variability of the duration of the dominating tasks. Most of the pyruvateKinase_MN.mdrun_pc_all (97%) have a duration between 215 - 229 minutes. There are a few of these tasks (3%) that take around 275 minutes. However, this almost homogeneous behaviour is not observed in all the tasks. For example, for the pyruvateKinase_MN.mdrun_pc_cpt task, we observe three different behaviours: there are 25% of the tasks that take 37.5*s* in average; 50% of the tasks that take 65.3*s* on average, and 25% of the tasks that take around 279.7*s* in average.

Figure 10 shows an histogram of the duration of the task pyruvateKinase_MN.mdrun_pc_cpt where the triple behaviour mentioned above can be observed. This triple behaviour of the task has an explanation in the code: for each mutation, the task is executed in 8 phases with different parameters and a different number of steps (nsteps). For two of the phases, nsteps is 5,000; for 4 of the phases, nsteps is 10,000 and finally, for two other phases, nsteps is 50,000.

**Fig. 10.**
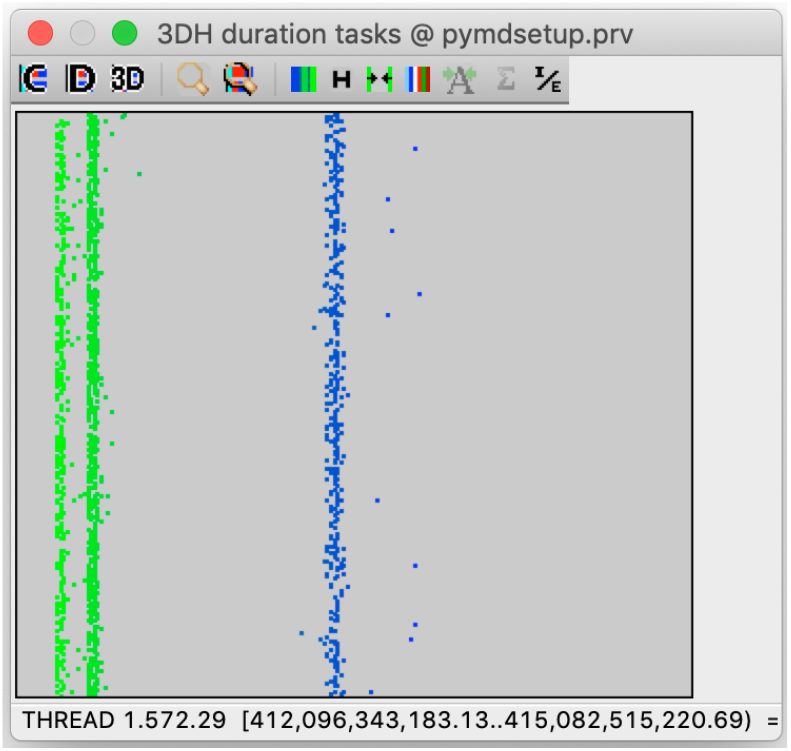
Paraver histogram showing the duration of the pyruvateKinase_MN.mdrun_pc_cpt task

The task pyruvateKinase_MN.mdrun_pc also shows variability: in this case, half of the tasks (200) last on average 124.54*s* and the other half last 505.89*s* on average. In this case, the slower tasks (that also show higher variability) are executed after the faster ones.

## V. Related Work

The use of computational workflows has become ubiquitous for data analytics in the field of bioinformatics since the last decade. In the literature, more than 200 workflows systems [14] can be found, targeting specific scientific domains, different execution models and usability approaches. Workflow systems can be classified according to the model used to define the tasks and the data dependencies and to the characteristics of the engine that executes the workflow on the computing platform. With relation to the tasks definition features, some frameworks allow to explicitly define the workflow through a recipe file or a graphical interface while others permit the users to program their applications and let the runtime build a dependency graph from the user code. Another relevant characteristic for the classification of these frameworks is the level of integration with the different computing platforms as distributed environments (such as grids, clouds, and clusters), and HPC systems with multi-core architectures and accelerators (such as GPGPUs).

Amongst all these tools, in this paper we focus on the features that are more convenient for the orchestration of molecular dynamics simulations, taking into account interoperability across a variety of software and hardware environments, scalability, and reproducibility. In particular, we consider HPC-focused workflow managers that can compose and run workflows with advanced features as elasticity, adaptability, and fault tolerance.

Taverna [15], [16], Kepler [17], [18], Galaxy [19], [20] are well known graphical environments for the composition of workflows that can be stored and shared with other users of the community and that are typically executed on supercomputers, grids, or cloud environments. KNIME [21] Analytics Platform also includes a graphical interface and supports external frameworks like Keras and Spark.

More bioinformatics specific environments have been recently developed. Crossbow [22] is a Python-based toolkit for workflow construction and execution, aimed mainly at Crossbow clusters but more generally at distributed computing environments. It provides an easy entry to cloud-based computing for biomolecular simulation scientists. Crossbow shares many of its design aspects with Parsl [23]. It provides tools to wrap Python functions and external applications (e.g., legacy MD simulation codes), in such a way that they can be combined into workflows using a task-based paradigm. Crossbow uses Dask [24] Distributed as the task scheduling and execution layer.

RADICAL-Cybertools [25] enable the execution of ensemble-based applications on a variety of high performance computing infrastructures. An increasing number of scientific domains are adopting and benefiting from ensemble-based applications. Most notably, MD simulations are nowadays executed as many parallel jobs of ns-length simulations rather than a single, long, and very large MPI job. AdaptiveMD [26] is a Python package designed to create HPC-scale workflows (parallel tasks) for adaptive sampling of biomolecular MD simulations. AdaptiveMD is designed as a distributed application that can be launched from a laptop or directly on an HPC resource and asynchronously automate the workflow creation and execution. Multiple adaptive sampling algorithms are fully automated with minimal user input, while advanced users can easily make modifications to workflow parameters and logic through the Python API. Runtime adaptations include the use of interim data as task properties such as analysis types or parameters, and workload properties such as task count or convergence criteria. To provide robust workflow management, AdaptiveMD is also integrated with the RADICAL Cybertools stack, which significantly enhances the runtime error detection and correction functionality, but has a much higher installation and configuration overhead.

The solution described in this proposal advances the mentioned approaches in the move to developing robust and scalable scientific workflow without the requirement of deep programming knowledge on the users. The adoption of Py-COMPSs provides powerful features which simplify the development and executions of complex workflows combining several types of heterogeneous tasks running in parallel on thousands of computing cores. Graphical workflow systems like Galaxy and KNIME have generally limited support for using HPC and HTC compute infrastructure in combination with high-performance codes like GROMACS, while our solution provides a solid solution for the execution of applications on a lot of computing backends withou the need of adapting the code to a specific one.

## VI. Conclusions

The paper has presented a work to reduce the gap between biomolecular research and the high performance computing world. The motivation for this work comes from the analysis, performed in the context of the BioExcel project, of the current situation around the execution of biomolecular workflows in supercomputing facilities.

The proposal has been developed around two pillars: usability and efficiency. A library of platform agnostic building blocks for molecular dynamics has been used to address the usability requirement, and pipelines made of these blocks have been defined adopting a task based approach that has a minimal impact on the code and that enables parallelization at execution time. Efficiency in the execution of the resulting workflows on supercomputing premises has been achieved through the parallel execution of different mutations using the COMPSs programming framework. The results of the runs, on up to 38,400 cores of the MareNostrum IV supercomputer, demonstrate that the workflows can be easily scaled to a large number of nodes and that the optimal configuration of the execution parameters can be obtained without modifying the user code. A complete analysis of the executions also proves that the workflow manager selected in our approach allocates the resources for the different types of tasks in the proper way with an almost 100% of utilization of the pool.

Future work is based on a well defined roadmap whose tasks will be performed in the BioExcel-2 project. The first topic is focused on the development of more complex workflows, extending the set of building blocks library, to exploit the power of the current supercomputer generation approaching the exascale paradigm. In connection with this topic, another relevant part in the roadmap is the introduction of data analytics in the workflows, in particular, the adoption of High Performance Data Analytics (HPDA) coupled to High Performance Computing (HPC). The use of HPC parallel processing to run powerful data analysis software tools opens the possibility to examine large (even massive) datasets within a reasonable time. A set of HPDA building blocks will be developed, starting with a recently developed library for distributed computing integrated on top of the PyCOMPSs framework and focused on machine learning. The last part of the work plan addresses the enhancement of the programming framework interfaces to deal with the requirement of porting the workflows to (pre)exascale facilities. These extensions allow to express dynamicity and support the description of iterative constructions such as conditional loops, still offering the full expressiveness of the programming language for complex algorithms, like optimization searches.

## ACKNOWLEDGMENT

This work has been supported by the Spanish Government (SEV2015-0493), by the Spanish Ministry of Science and Innovation (contract TIN2015-65316-P), by the Generalitat de Catalunya (contract 2014-SGR-1051), and by the European Commission through BioExcel Center of Excellence (Horizon 2020 Framework program) under contracts 823830, and 675728. Cristian Ramon-Cortes predoctoral contract is financed by the Ministry of Economy and Competitiveness under the contract BES-2016-076791.

## Notes

### Competing Interest Statement

The authors have declared no competing interest.

